# Light regulates widespread plant alternative polyadenylation through the chloroplast

**DOI:** 10.1101/2024.05.08.593009

**Authors:** M. Guillermina Kubaczka, Micaela A. Godoy Herz, Wei-Chun Chen, Dinghai Zheng, Ezequiel Petrillo, Bin Tian, Alberto R. Kornblihtt

## Abstract

Transcription of eukaryotic protein-coding genes generates immature mRNAs that are subjected to a series of processing events, including capping, splicing, cleavage and polyadenylation (CPA) and chemical modifications of bases. Alternative polyadenylation (APA) greatly contributes to mRNA diversity in the cell. By determining the length of the 3’ untranslated region, APA generates transcripts with different regulatory elements, such as miRNA and RBP binding sites, which can influence mRNA stability, turnover and translation. In the model plant *Arabidopsis thaliana*, APA is involved in the control of seed dormancy and flowering. In view of the physiological importance of APA in plants, we decided to investigate the effects of light/dark conditions and compare the underlying mechanisms to those elucidated for alternative splicing (AS). We found that light controls APA in approximately 30% of *Arabidopsis* genes. Similar to AS, the effect of light on APA requires functional chloroplasts, is not affected in mutants of the phytochrome and cryptochrome photoreceptor pathways and is observed in roots only when the communication with the photosynthetic tissues is not interrupted. Furthermore, mitochondrial activity is necessary for the effect of light in roots but not in shoots. However, unlike AS, coupling with transcriptional elongation does not seem to be involved since light-dependent APA regulation is neither abolished in mutants of the TFIIS transcript elongation factor nor universally affected by chromatin relaxation caused by the histone deacetylase inhibition. Instead, regulation seems to be linked to light-elicited changes in the abundance of constitutive CPA factors, also mediated by the chloroplast.

## Introduction

Transcription of eukaryotic protein-coding genes generates immature mRNAs that are subjected to a series of co-transcriptional processing events, including capping, splicing, cleavage and polyadenylation (CPA) and chemical modifications of bases. CPA is responsible for cleavage of the nascent RNA and addition of a poly(A) tail to cleaved RNA. Most of the core CPA factors found in yeast and mammalian cells, about 20 in total, have homologous counterparts in the plant *Arabidopsis thaliana*, in particular within the complexes CPSF (cleavage and polyadenylation specificity factor), CstF (cleavage stimulation factor) and CFII (cleavage factor II). Equivalent subunits of mammalian CFI (cleavage factor I), however, have not been identified in plants (1). In all studied organisms, core CPA factors act together with the poly(A) polymerase and other RNA-binding proteins to yield poly(A)-tailed mature mRNAs. A key step in this process is the recognition of RNA motifs that define the polyA site (PAS). The AAUAAA hexamer is the most prominent PAS motif in metazoans, which is recognized by CPSF. However, this sequence has only been reported in approximately 10% of the *Arabidopsis* transcripts, which opens the yet-unsolved question of how the PAS is specified in plants (1).

Similar to alternative splicing (AS), alternative polyadenylation (APA) greatly contributes to mRNA diversity in the cell (2). By affecting the length and sequence of the 3’ untranslated region (3’UTR), APA generates transcripts with different architectures of target sites for microRNAs, which affects mRNA turnover and translation. When coupled with AS, APA can also alter C-terminal segments of the polypeptides encoded by a single gene. For example, this mechanism is involved in the control of seed dormancy in plants. The *Arabidopsis DOG1* (delay of germination 1) gene gives rise to two mRNA isoforms that are generated by the use of proximal and distal (with respect to the promoter) PASs, respectively (3). The short and long mRNAs encode polypeptides of 30 and 32 kDa respectively, differing in a non-conserved C-terminal stretch. However, only the protein encoded by the short mRNA is functionally active in promoting dormancy. Mutations in CPA factors that lead to reduced use of the proximal PAS cause weakened seed dormancy (3). APA of non-coding RNAs also has fundamental biological roles in plants. The best studied example is the role of the long non-coding RNA *COOLAIR* in the control of flowering in *Arabidopsis. COOLAIR* is an antisense transcript of the flowering locus C gene (*FLC*) that displays two APA sites. When the distal site is used, a transcription-permissive histone mark is deployed along the FLC gene which allows for FLC expression and repression of flowering. Instead, if the proximal site is used, the short isoform of *COOLAIR* base-pairs with complementary *FLC* sequences, forming a so-called R-loop, which promotes the deployment of a transcription-repressive histone mark, which subsequently inhibits *FLC* expression and promotes flowering (4, 5).

We have previously shown that light regulates *Arabidopsis* AS through the chloroplast. We demonstrated that the photosynthetic electronic transport chain initiates a chloroplast retrograde signaling that regulates nuclear splicing (6). Later we found that light promotes RNAPII elongation while in darkness elongation is lower (7). These changes in transcriptional elongation are causative of the observed changes in AS, indicating that the chloroplast control responds to the kinetic coupling between transcription and RNA processing found in mammalian cells (for reviews see references 8 and 9), and providing unique evidence that coupling is important for a whole organism to respond to environmental signals.

In view of the physiological importance of APA in plants, we decided to investigate the effects of light/dark conditions and the underlying mechanisms and to compare them to the ones elucidated for AS. We report here that light deeply affects *Arabidopsis* 3’UTR APA in 28.7% (3,467) of 12,070 genes assessed by genome-wide analyses. Three gene groups are clearly defined. Those in which light promotes preferred usage of proximal PASs, those in which distal PASs are preferred and those in which light has not effect on APA at all. Similar to AS, the effect of light on APA requires functional chloroplasts, is not affected in mutants of the phytochrome and cryptochrome photoreceptor pathways, is observed in roots when the communication with the photosynthetic tissues is not interrupted. Furthermore, like in AS, mitochondrial activity is necessary for the APA effect of light in roots but not in shoots (10). However, unlike AS, coupling with transcriptional elongation does not seem to be involved in changes in APA triggered by light. Alternatively, regulation seems to be linked to light-elicited changes in the abundance of constitutive CPA factors, also mediated by the chloroplast.

## Results

### Arabidopsis genes display substantial 3’UTR APA isoform expression

In order to investigate the role of light/dark exposure in *Arabidopsis* APA, we applied a light regime similar to the one used in our previous AS studies (6, 7, 10). Briefly, seedlings were grown for two weeks in constant white light to minimize interference from the circadian clock and then transferred to light or dark conditions for different time periods (Fig. 1A). To obtain APA regulation information genome-wide, we subjected total RNA to RNA sequencing by using the 3’ region extraction and deep sequencing (3’READS) method (11, detailed in Materials and Methods). We generated ∼82.5 million PAS-supporting reads from the two sample groups (Suppl. Table 1). After clustering of adjacent cleavage sites that resulted from heterogenous cleavage for each PAS (Suppl. Fig. 1A, see Materials and Methods for detail), we identified 160,881 PASs in the *Arabidopsis* genome. Consistent with previous studies (12, 13), plant PASs identified by 3’READS are surrounded with A/T-rich sequences, with a prominent T-rich peak around –10 nt and a modest A-rich peak around –20 nt (cleavage site is set to position 0, Suppl. Fig. 1B). By analyzing hexamers in neighboring regions of the PAS, we found substantially enriched TA-rich motifs surrounding *Arabidopsis* PASs (Suppl. Fig. 1C), which is similar to PASs in *S. cerevisiae* but distinct from those in metazoans (14–17).

**Figure 1.**
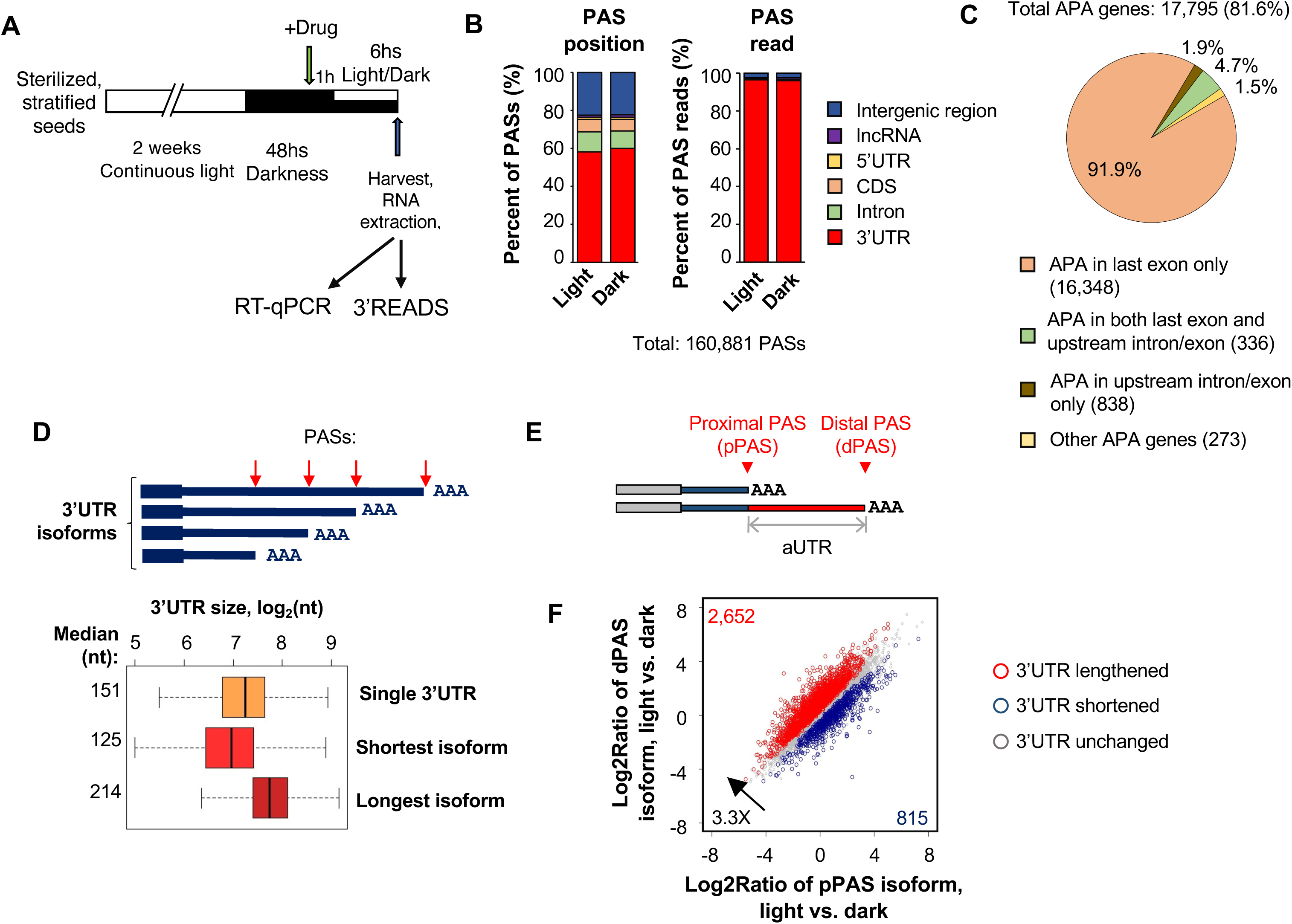
Light/dark conditions elicit widespread APA changes in *Arabidopsis thaliana*. **A**. Protocol scheme of the light/dark regime used in this study. Total RNA from seedlings was subject to 3’READS for gene expression and APA analyses. **B.** Distribution of identified PASs (left) and PAS reads (right) in different regions of the plant genome. **C.** Different types of APA genes identified in this study. **D.** Top, schematic of 3’UTR APA isoform; bottom, 3’UTR sizes for mRNAs of genes without 3’UTR APA (single 3’UTR) and mRNAs with the longest or shortest 3’UTRs of genes with 3’UTR APA. The median value for each group is indicated. **E.** Diagram showing 3′UTR APA analysis. The two most abundant APA isoforms per gene were selected for comparison, which are named proximal PAS (pPAS) and distal PAS (dPAS) isoforms, respectively. The distance between the two PASs is considered alternative 3′UTR (aUTR). **F.** Scatterplot showing genes with pPAS and dPAS isoform abundance differences between light– and dark-treated seedlings. Results represent analyses of 12,873 genes in three biological replicates. Genes with significantly (FDR < 0.05, DEXseq analysis) higher or lower abundance of pPAS isoforms in light vs. dark conditions are shown in blue and red respectively.

Overall, 59% of all PASs were mapped to the last exon of mRNA genes, mostly in 3’UTRs. In addition, 10% were in introns, 6% in coding exons and 22% in intergenic regions (Fig. 1B, left). Notably, the vast majority of the PAS-supporting reads (96%) were mapped to the last exon (Fig. 1B, right), indicating that transcripts using last exon PASs are much more abundant than those using PASs in other regions.

Using transcript abundance of 5% as the cutoff to call an isoform (Suppl. Fig. 2A), we found that 81.6% of plant mRNA genes display APA in our samples. On average, an APA gene expresses 2.9 APA isoforms (Suppl. Fig. 2B). Of all APA genes, 91.9% have APA sites exclusively in the last exon and 4.7% have APA sites in both the last exon and upstream introns/exons (Fig. 1C). Therefore, most plant genes display 3’UTR APA through alternative usage of PASs in the last exon.

For genes that do not display APA, the median value of their 3’UTR size is 151 nt (Fig. 1D). By contrast, for genes that expression 3’UTR APA isoforms, the median values for the shortest 3’UTR and the longest 3’UTR are 125 nt and 214 nt, respectively (Fig. 1D). The alternative 3’UTR size has a median value of 78 nt across genes (Suppl. Fig. 2C).

Therefore, 3’UTR APA in plants can potentially alter a substantial portion (30-60%) of the 3’UTR sequence, impacting post-transcriptional control of gene expression. We also found that the longest 3’UTR isoform is typically expressed at a higher level than the shortest 3’UTR isoform (Suppl. Fig. 2D). In addition, the PASs of the shortest and longest isoforms are surrounded with distinct mRNA motifs (Suppl. Fig. 2E), with the AATAAA motif highly enriched for the last PAS (longest 3’UTR isoform) and TGT motif for the first PAS (shortest 3’UTR isoform).

### Light/dark switch elicits widespread APA isoform changes

We next examined APA isoform changes in light vs. dark conditions. Because most genes display 3’UTR APA, we focused on the top two most expressed 3’UTR APA isoforms for each gene. For simplicity, they are named proximal PAS (pPAS) isoform and distal PAS (dPAS) isoform, respectively (Fig. 1E). We calculated their relative expression (RE, dPAS vs. pPAS) in each sample group and then compared RE between the groups (light vs. dark), yielding the value RED (relative expression difference).

Using p-value cutoff of 0.05 (DEXSeq), we identified 3,467 genes that showed significant differences in APA isoform expression levels between light and dark conditions (Fig. 1F). Strikingly, genes showing 3’UTR lengthening, i.e., upregulation of dPAS isoform relative to pPAS isoform, outnumbered those showing 3’UTR shortening by 3.3-fold (Fig. 1F). By using Gene Ontology analysis, we found that genes showing 3’UTR lengthening tend to have functions in various metabolic pathways, such as ATP, cellular amide, hexose, etc. (Suppl. Table 2).

Representative genes showing 3’UTR lengthening, 3’UTR shortening or no change are shown in Suppl. Fig. 3. *HTA9* and *RKH* are two paradigmatic examples of 3’UTR lengthening and 3’UTR shortening genes, respectively (Fig. 2A). Using PCR primers targeting alternative 3’UTR sequences of their isoforms and upstream common regions (Suppl. Table 3), we validated our 3’READS data by RT-qPCR.

**Figure 2.**
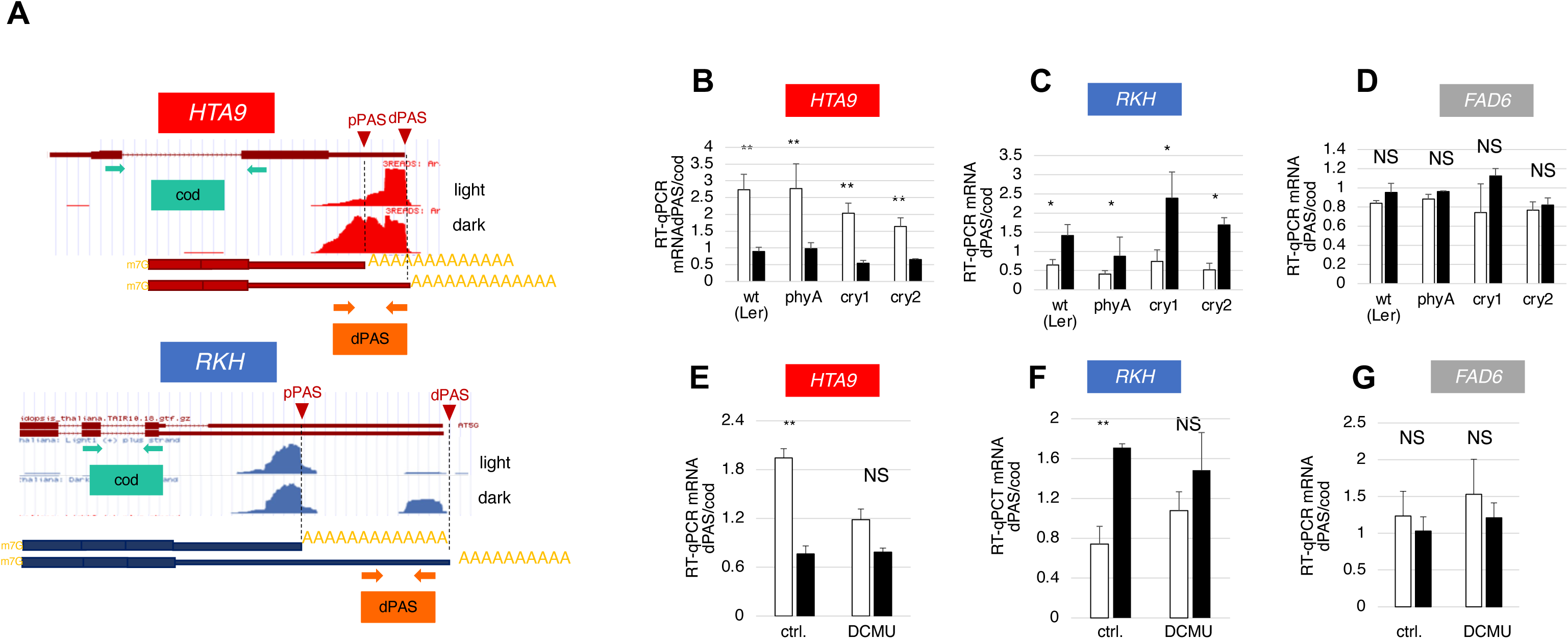
The light effect on APA is sensed by the chloroplast (genetic and biochemical evidence). **A**. 3’READS data for two *Arabidopsis* genes with opposite APA changes. Top: a representative event [*HTA9* gene (AT1G52740), reads in red] with higher usage of dPAS in the light. Bottom: a representative event [*RKH* gene (AT5G15270), reads in blue] with higher usage of pPAS in the light. For each APA event two pairs of primers were designed to validate the APA changes using RT-qPCR: amplicon dPAS (dark orange) only exists if the dPAS is used (long isoform); amplicon cod is common to all isoforms in the upstream coding region. Changes in APA are quantified as ratios of dPAS/cod amplicons relative mRNA expression levels for every gene. **B-D.** Light is not sensed by photoreceptors. APA response to light/dark in different *Arabidopsis* phytochrome and cryptochrome mutant genotypes in a Landsberg erecta background (wt, Ler). Three selected genes are shown: *HTA9*, whose APA events increases its dPAS usage in the light (A), *RKH*, whose APA event diminishes its dPAS usage in the light (B) and *FAD6*, whose APA event is not affected by the light/dark conditions, and serves as a negative control. **E-G**. Effect of the photosynthetic electron transfer chain inhibitor DCMU on the light/dark effect on APA events of the *HTA9* (**E**), *RKH* (**F**) and *FAD6* (**G**) genes. Seedlings were grown in constant light, transferred to darkness for 48 hr. and then treated with 20 µM DCMU during a 6-hr. light/dark further incubation. RT-qPCR experiments were quantified with n ≥ 3, where n = ∼ 25-30 *Arabidopsis* seedlings growing in one Petri dish. White and black bars represent light and dark treatments respectively. Changes considered significant show differences with a p value < 0.05 (two-tailed Student’s t test). *** = p < 0.001; ** = p < 0.01; *= p < 0.05; NS (not significant) = p > 0.05.

Our 3’READS data also detected widespread gene expression changes in light/dark conditions (Suppl. Fig. 4A). We found that there is a significant bias for upregulated genes having 3’UTR shortening and downregulated genes having 3’UTR lengthening (P = 0.002, Fisher’s exact test, Suppl. Fig. 4B). This result indicates that 3’UTR APA is relevant to gene expression changes.

### The chloroplast is the sensor for the light effect on APA

Next we wanted to address whether signaling through photoreceptors was involved in APA regulation in response to light. We exposed seedlings of *Arabidopsis* mutants for the red/far red-light photoreceptor phytochrome A (phyA-201, 18) and for the blue light photoreceptor cryptochrome cry1 (cry1-1, 19) and cry2 (fha-1, 20) to our light regime protocol (Fig. 2B-D), using the wild type Landsberg erecta (Ler) background as control. APA was assessed by RT-qPCR in the *HTA9* and *RKH* genes, as examples of 3’UTR lengthening and shortening by light respectively (Figs. 2B and 2C), and the *FAD6* gene (Fig. 2D) as a gene with no change in APA. Both types of mutants behaved similarly to wild type seedlings for all three genes, which refutes the notion that photoreceptors play a role in on APA.

Because retrograde signals from the chloroplast have been shown to modulate nuclear gene expression (6, 10, 21), we reasoned that chloroplast functions might be involved in light-elicited APA regulation. To this end, we used the herbicide DCMU [3– (3,4-dichlophenyl)-1,1-dimethylurea] (22) that blocks the photosynthetic electron transport from photosystem II to the plastoquinones. Interestingly, DCMU inhibited the effect of light on APA in both *HTA9* (Fig. 2E) and *RKH* (Fig. 2F) genes but had no effect on *FAD6* (Fig. 2G), suggesting that chloroplast function is necessary for modulating APA in response to light.

### The light effect on APA is sensed by the photosynthetic tissues

To obtain further evidence of the involvement of the chloroplast in APA we performed dissection experiments. Because roots have no chloroplasts, we reasoned that if seedlings were cut to separate roots from green photosynthetic tissues (shoots) (Figs. 3A and 3B), APA differences upon light/dark treatments should only be observed in the green tissues. Similarly to what happens with AS (6), in the case of the *HTA9* gene the effect was observed both in dissected leaves and roots (Figs. 3C, left) when dissection was performed 6 hr after light/dark treatment. This led us to perform a dissection experiment in which shoots were separated from roots before the treatment (Fig. 3C, right). In these conditions dissected shoots retain the same light response for *HTA9* APA as undissected seedlings, but light has no effect on APA the in dissected roots. In the case of *RKH* APA, the light effect was only observed in shoots independently of whether dissection was performed after or before treatment (Fig. 3D). The negative control *FAD6* was unresponsive in neither shoot nor roots in both dissection protocols (Fig. 3E). These results strongly reinforce the evidence that light modulates APA through the chloroplast. The *HTA9* dissection experiments suggest that a signal generated by light in the green tissues can move to the roots to modulate APA in a similar way as in the green parts of the plant.

**Figure 3.**
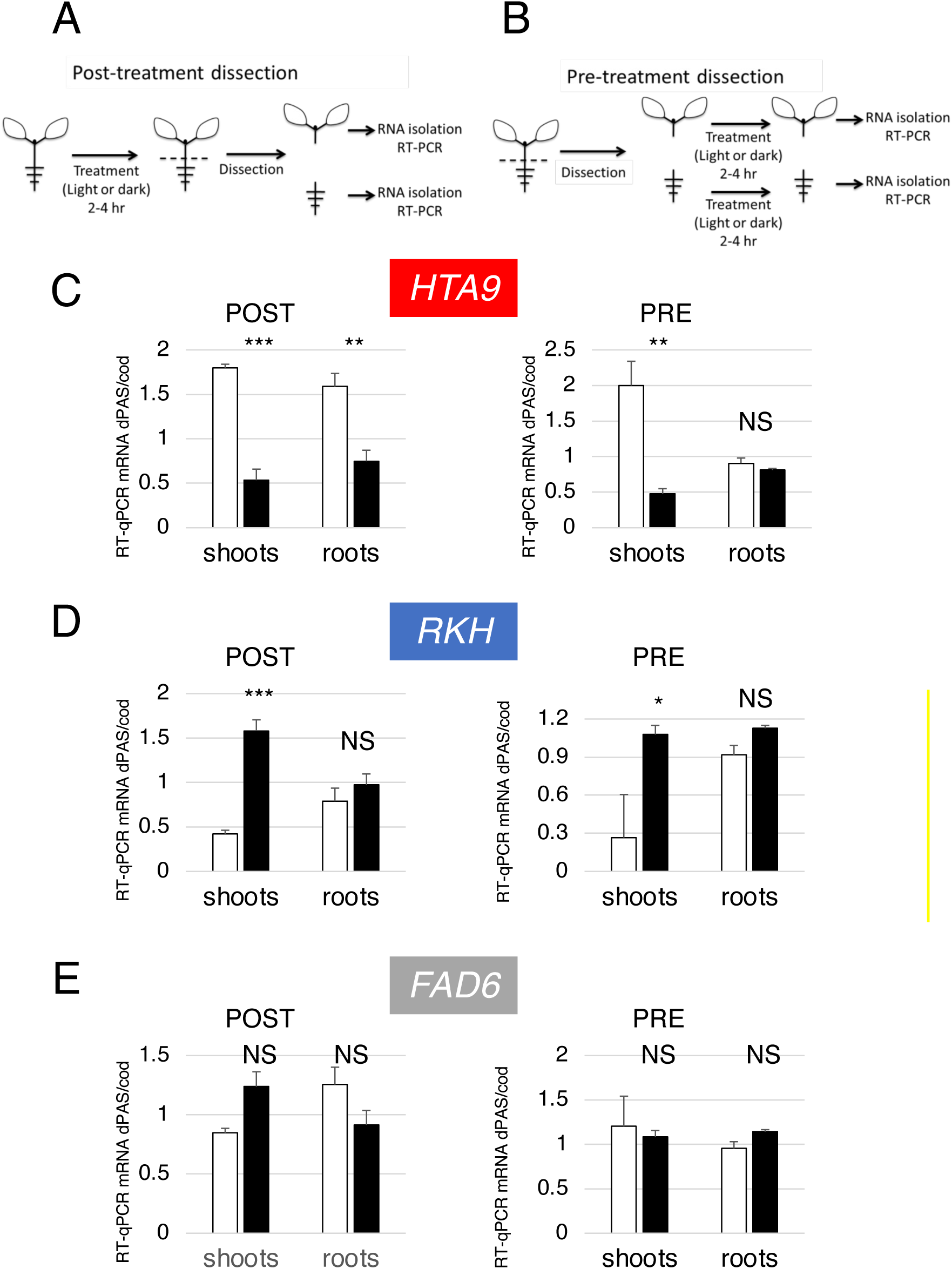
The light effect on APA is sensed by the photosynthetic tissues (anatomical evidence). **A-B**. Schemes for dissections of *Arabidopsis* seedlings performed after (post-) or before (pre-) the light/dark treatment. **C-E.** APA isoform analysis in green tissue (shoots) and in the roots of post-(left) and pre-dissection (right) light/dark treatments of the *HTA9* (C), *RKH* (D) and *FAD6* (E) genes. Bar colors and RT-qPCR conditions were as in Figure 2.

### Sugars and mitochondrial activity modulate the light effect on *HTA9* APA in the roots

In view if the evidence emerging from DCMU inhibition and dissection experiments, we decided to further investigate the signaling from shoots to roots. Sugars are the main photosynthates in terrestrial plants. These are generated in the green tissues and either metabolized in their cells or loaded into the phloem to feed non-photosynthetic tissues. In a previous study (10), it was shown that sucrose, the most important phloem-mobile sugar of *Arabidopsis* (23), is responsible of mediating AS changes in the roots triggered by light exposure of the shoots. More interestingly, externally applied sucrose solutions mimicked the effect of light on AS patterns in the root but had no major effect in the shoots. In agreement with AS results, incubation with 100 mM sucrose of seedlings subjected to the light/dark regime that were later dissected (Fig. 4A, POST) did not altered the light effect on *HTA9* APA in the shoots, but abolished it in the roots, in a way that mimics the effect of light in roots kept in the dark (note the height of the black bar that corresponds to roots treated with sucrose in Fig. 4A). In order to rule out any osmotic effect of sucrose we used equal concentrations of sorbitol as negative control. When dissection was performed before light/dark treatments (Fig. 4 B, PRE) incubation with sucrose had neither effect on the change in *HTA9* APA observed in the shoots nor on the lack of it observed in the roots.

**Figure 4.**
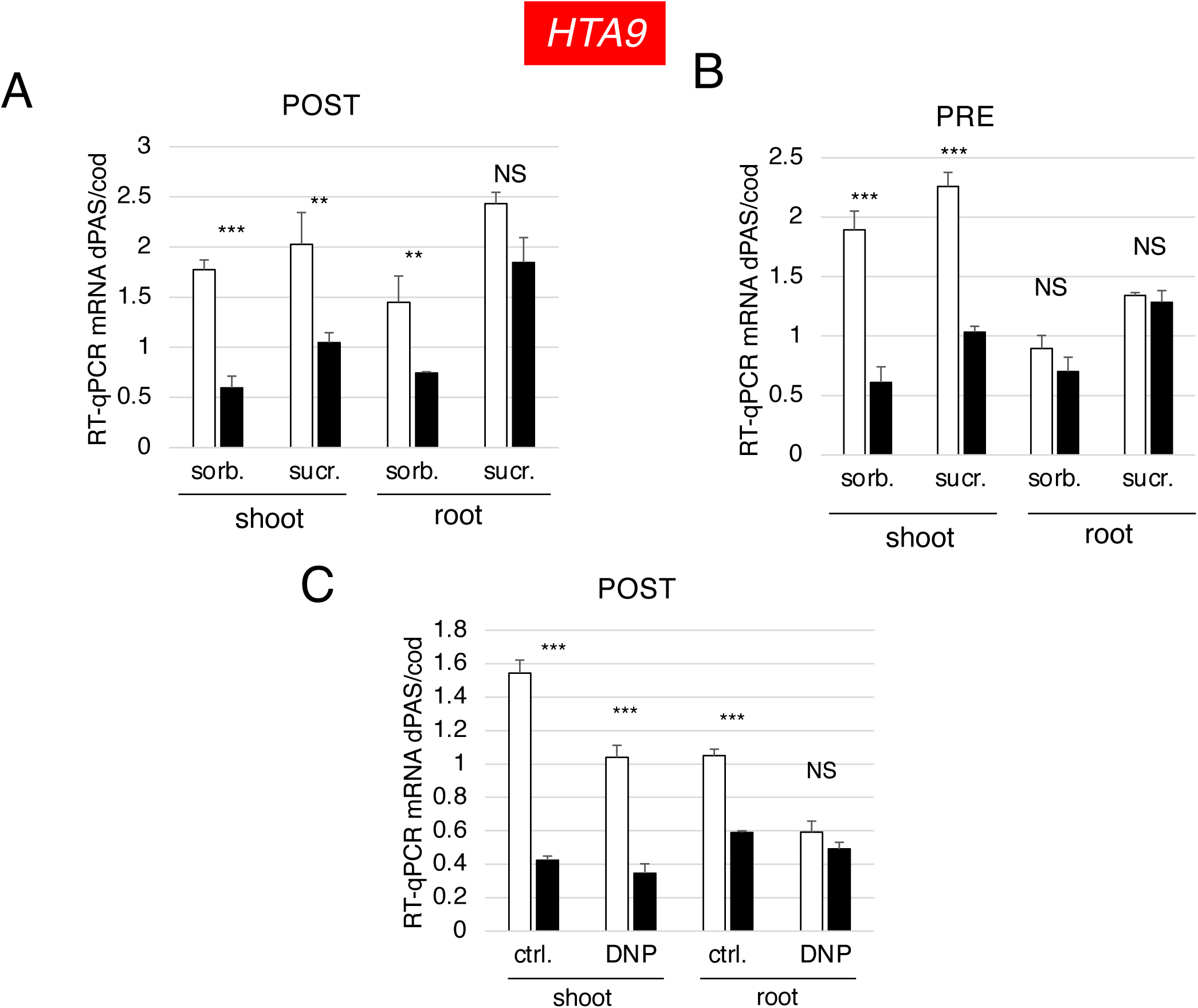
Sugars and mitochondrial activity modulate the light effect on APA in the roots. *HTA9* gene APA isoform analysis in green tissues (shoots) and roots of *Arabidopsis* seedlings dissected post-(**A** and **C**) and pre-(**B**) light/dark treatments. **A** and **B**. Incubations were performed with 100 mM sucrose or sorbitol (negative osmotic control) as indicated. C. Post-treatment excision protocol. Incubations were performed with 20 µM of the mitochondrial uncoupler dinitrophenol (DNP) or vehicle (ctrl.) as indicated. Bar colors and RT-qPCR conditions were as in Figure 2.

Sugars have dual roles in plants, serving both as metabolic fuel and as signaling molecules (24). The signaling pathway in which sugars activate target of rapamycin (TOR) kinase and subsequently gene expression has been involved in the regulation of AS by sugars in *Arabidopsis* roots (10). Xiong et al. (25) revealed that glucose activation of TOR kinase in *Arabidopsis* meristems depended on glycolysis-mitochondria-mediated energy. This is consistent with findings that the respiratory chain/oxidative phosphorylation uncoupler 2,4-dinitrophenol (DNP) suppresses mammalian target of rapamycin (mTOR) activation in brain (26). Since DNP was also shown to abolish changes in root AS induced by light exposure of the green tissues (10), we investigated the effects of DNP on APA. In order to be able to assess DNP effects in the roots, experiments were carried out in the POST protocol, i.e., where excision was performed after the light/dark or drug treatments. Fig. 4C shows that DNP does not suppress the light effect on APA in the shoots, but completely abolishes the conspicuous light effect in the roots, in a way that imitates darkness. Control experiments with the APA unresponsive FAD6 gene are shown in Suppl. Fig. 5.

### Unlike AS, light control of *HTA9* APA is not affected by transcriptional elongation

We used two experimental approaches to evaluate if the effect of light on *HTA9* APA was linked to changes in transcriptional elongation according to the kinetic coupling mechanism. To inhibit transcriptional elongation rate, we assessed an *Arabidopsis* mutant of the TFIIS transcription elongation factor. TFIIS is a factor required for RNAPII processivity that stimulates RNAPII to reassume elongation after pausing (27, 28). The *tfiis* mutant was achieved by replacing the key amino acids D290 and E291 of the acidic loop responsible for TFIIS stimulatory activity by alanines (28) as described in yeast (29), giving rise to a dominant negative phenotype showing a range of developmental defects, such as defective growth and serrated leaves. We have previously reported that in the *tfiis* mutant, transcript elongation is reduced and the change in *AtRS31* AS induced by light is abolished, mimicking the effects of darkness (7). While this is confirmed as a positive control in Fig. 5A, the effect of light on *HTA9* APA is not affected in the *tfiis* mutant (Fig. 5C). The negative control *FAD6* behaves similarly (Fig. 5E). On the activation side, we explored the use of a histone deacetylase inhibitor (HDAC) trichostatin A (TSA). By promoting higher histone acetylation and chromatin opening, HDACs have been proved to promote elongation in animal (30, 31) and plant cells (7), and in the latter, to mimic the chloroplast-mediated light effect on *AtRS31* AS. Again, while this is confirmed as positive control in Fig. 5B, the effect of light on *HTA9* APA is not affected by TSA treatment (Fig. 5D) and the negative control *FAD6* behaves similarly (Fig. 5F).

**Figure 5.**
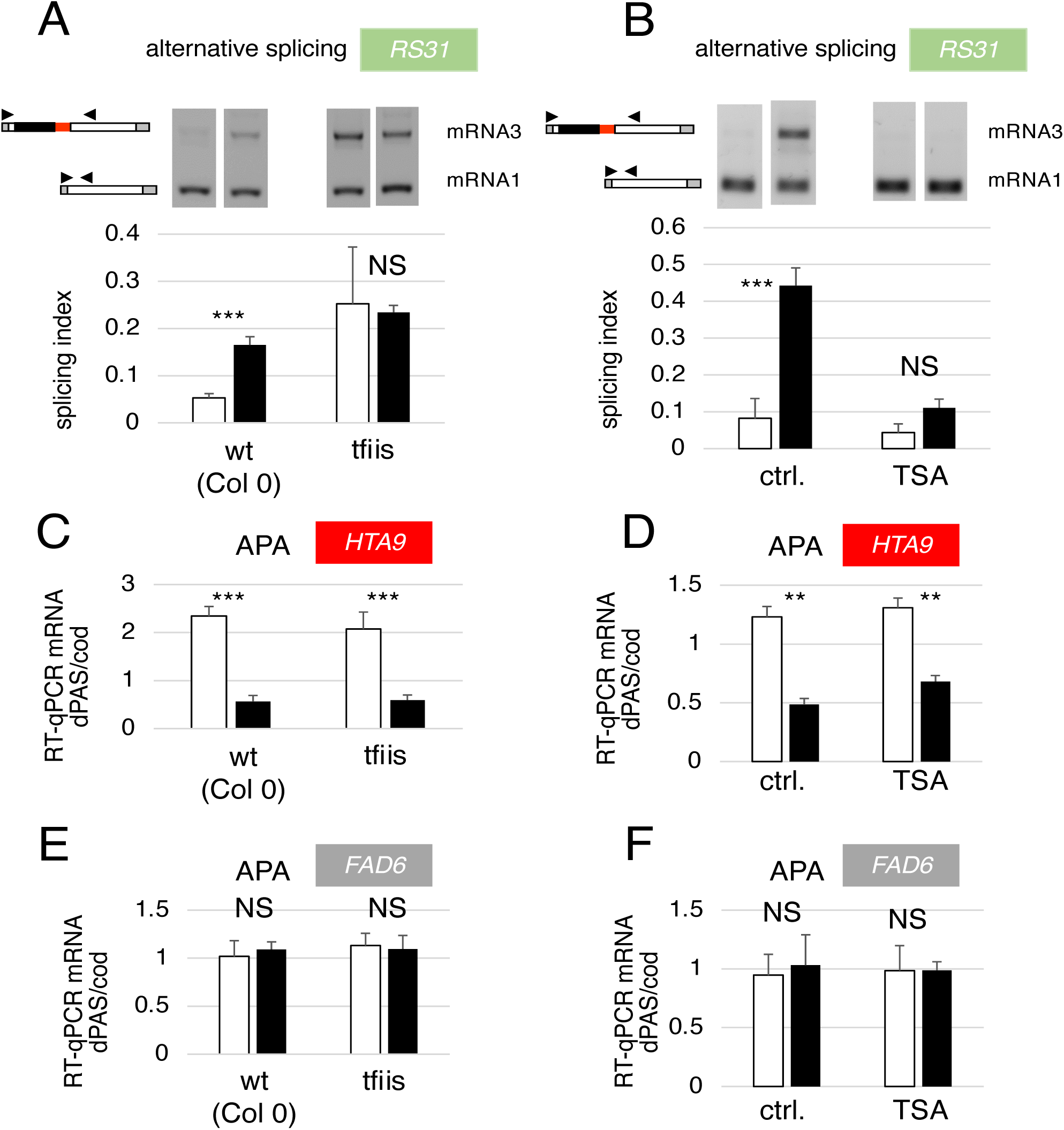
AS and APA respond differently to factors affecting transcript elongation. Effects of genetic disruption (tfiis mutant) of the transcription elongation factor TFIIS (A, C and E) and of treatment with the histone deacetylase inhibitor TSA (B, D and F) on AS of the *Arabidopsis RS31* gene (A and B) and on APA of the *HTA9* (C and D) and *FAD6* (E and F) genes. Bar colors and RT-qPCR conditions were as in Figure 2.

### Changes in CPA factor mRNA abundance

In view of the fact that the light effect on APA does not seem to follow the kinetic coupling mechanism, we decided to investigate if the changes in APA in the light could be explained by changes in the expression of CPA factors measured at the mRNA level. We assessed three subunits of CPSF and two of CstF in shoots and roots obtained from excision performed after the light/dark treatment. All three CPSF subunits (100, 160 and 30) exhibited reduction in their steady state mRNA levels in the dark both in the shoots and in the roots (Fig. 6A). The effect is smaller in the roots (note smaller values for the white bars) suggesting that a signal generated in the green tissues reaches the roots but is not as effective as the signal in the shoots. In the case of CstF, while no changes in mRNA levels were observed for CstF64 neither in the shoots nor in the roots, CstF77 mRNA abundance drops in the dark in the shoots but is not affected in the roots (Fig. 6B). To investigate if, as shown for APA in Figs. 2E-G, the chloroplast is the sensor for the upregulation of CPSF100, 160 and 30 and CstF77 by light, we treated whole seedlings with the electron transport chain inhibitor DCMU. Consistently, the DCMU treatment abolished the light/dark effect on mRNA levels for these subunits (Figs. 6C and 6D).

**Figure 6.**
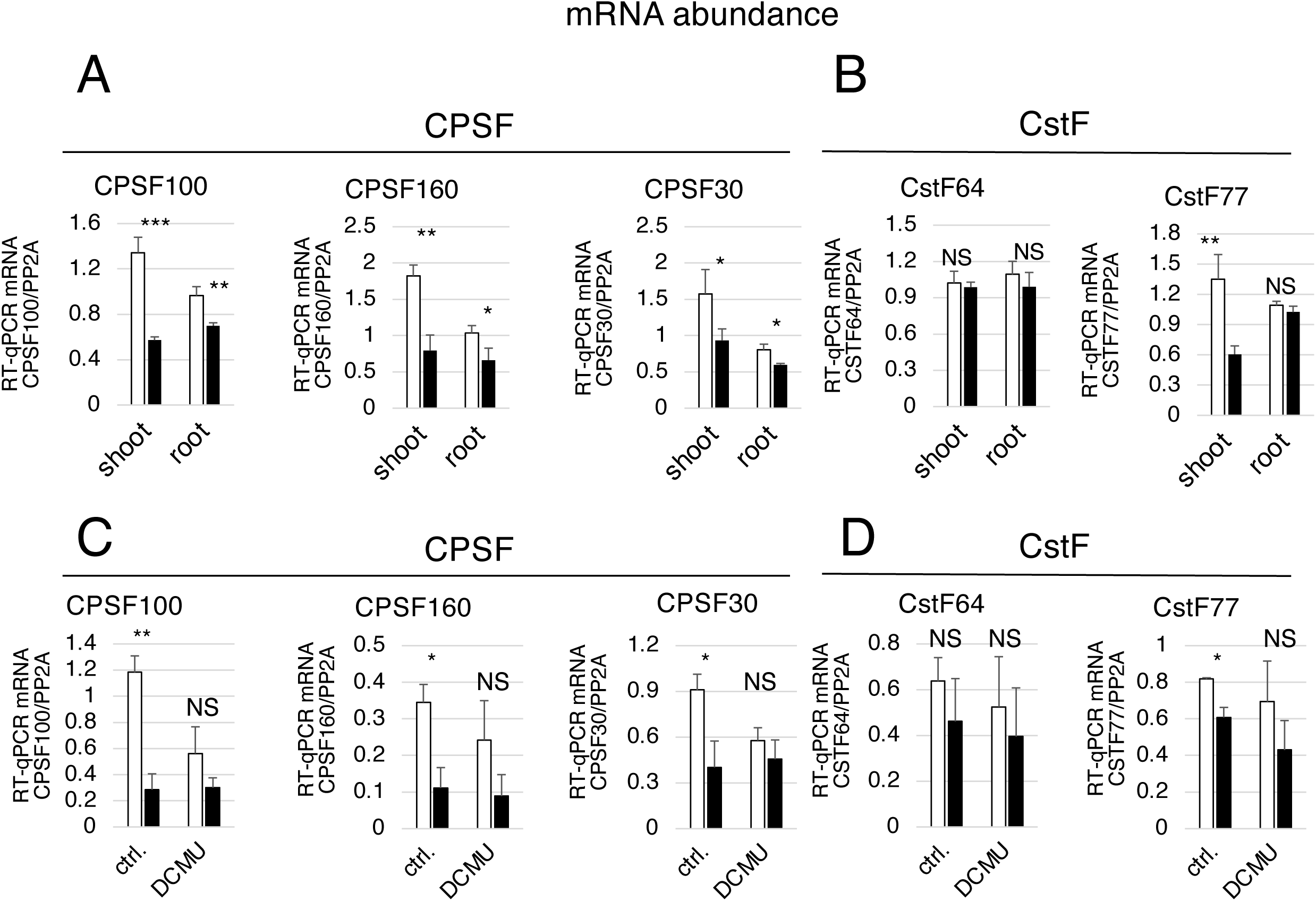
Light upregulates mRNA levels of cleavage/polyadenylation factors through the chloroplast. **A** and **B**. RT-qPCR quantification of mRNA levels encoding subunits of CPSF (**A**) and CstF (B) in *Arabidopsis* shoots and roots obtained in a post-light/dark treatment excision experiment. **C** and **D**. Effect of the photosynthetic electron transfer chain inhibitor DCMU on the light/dark effect on cleavage/polyadenylation factor mRNA levels in whole seedlings for CPSF (**C**) and CstF (**D**). Factor mRNA levels were relativized to mRNA levels of protein phosphatase 2A (PP2A). Bar colors and RT-qPCR conditions were as in Figure 2.

## Discussion

We report here an unforeseen, potent regulation of APA by light/dark conditions in plants. It should be noted that the effects here described occur in an intact living organism (*Arabidopsis* seedlings) and under a fundamental physiological external signal (light). Although this investigation was initially inspired by our previous work on the effect of light on plant AS (6, 7, 10), our findings indicate that, despite some similarities, APA regulation has fundamental differences with respect to that of AS. Light/dark conditions affect 3’UTR APA of a substantial number of *Arabidopsis* genes (approximately 30%), which highlights its global impact. In the regulated genes light promotes the usage of both dPAS (75% of genes) and pPAS (the remaining 25%). Because of the predominance of dPAS usage and the robustness of its response, we decided to use the APA event of a gene encoding a histone H2A variant (*HTA9*) as a model to investigate the mechanisms involved. An ideal event to study would have been APA in the long non-coding RNA COOLAIR (AT5G01675) that is key in the control of flowering (4, 5), but unfortunately, COOLAIR is not expressed in seedlings (32), the paradigm system of our previous work on splicing and the present work on polyadenylation.

Similar to the regulation of AS, the light effect on APA is not mediated by phytochrome nor cryptochrome photoreceptors but by the chloroplast. The correct functioning of the photosynthetic electron transport chain is necessary since its inhibition by DCMU abolishes the effect. This is consistent with the fact that the light effect is observed both in the green tissues and the roots as long as their connection is not interrupted by dissection. Since the roots have no functional chloroplasts, the absence of effect in isolated roots indicates that a signaling molecule must travel through the phloem from shoots to roots. We identify this molecule as the main photosynthate sucrose because it fully mimics the effect of light in the roots when seedlings are kept in the dark. Most interestingly, we found that root mitochondrial activity is necessary in the roots since the respiratory chain/oxidative phosphorylation uncoupler 2,4-dinitrophenol (DNP) abolishes the light effect fully mimicking the effect of darkness in the roots when seedlings are kept in the light. The opposite effects of sucrose and DNP allowed us to hypothesize that, like in AS regulation (10), the APA mechanism could involve sugar activation of TOR kinase, previously shown to depend on mitochondria-generated energy (25). Notably, 3’UTR size regulation through APA is increasingly associated with cell metabolism in mammalian cells (33). Whether plants and metazoans share similar mechanisms in energy-mediated APA regulation is to be examined in the future.

Two experiments of different nature indicate that, unlike AS, the effect on APA is not linked to the control of RNAPII elongation. The light effect on APA is neither abolished in a mutant of the transcription elongation factor TFIIS nor in seedling treated with the histone deacetylase inhibitor TSA, previously shown to cause higher transcript elongation due to histone acetylation and chromatin relaxation. Important controls show that both the TFIIS mutant and TSA treatment suppress the light effect on AS, the former mimicking darkness and the latter mimicking light as published (7). In search for an alternative mechanism we wondered if light would be affecting the expression of cleavage/polyadenylation factors. We found indeed that light upregulates mRNA levels for CPSF100, 160 and 30 and CstF77 and that this increase is suppressed by treatment with DCMU. In particular, *Arabidopsis* CPSF30 has been reported to interact with other polyadenylation factors like FIPS5, CPSF100, and CstF77 and can be considered as a central hub in the protein-protein interaction network of plant polyadenylation complex subunits (34–36). Indeed, *Arabidopsis* PAS selection, is determined by the presence or absence of this factor (37). The precise mechanism by which CPSF30 controls *Arabidopsis* APA remains elusive. However, the current model for canonical plant PA signaling involves three discreet cis-elements collaborating to the effective 3’ end formation of mRNAs: (i) the far upstream element (FUE), consisting of an extended U and G-rich region situated more than 50 nucleotides upstream from the PAS; (ii) the near-upstream element (NUE), a 6 to 10 nucleotide-A-rich region situated 10 to 30 nucleotides upstream from the PAS; and (iii) the cleavage element (CE), a U-rich region centered around the PAS. Although most functioning aspects of these elements have not yet been characterized, CPSF30 was shown to plays a role in the functioning of the NUE because PAS that are only used in the wild type, but not the CPSF30-deficient mutant, possess the characteristic A-rich NUE signature, while PAS used only in the CPSF30-deficient mutant lack this signature (37). In mammalian models, there is accumulated evidence that a variety of core CPA factors regulate APA (38–43). Moreover, a model was proposed in which the choice of PASs is dependent on the strengths of the cis-elements present in the PAS and the relative usage dependent on the competition between PASs for the available CPA factors (44). Upon loss or diminishment of core CPA factor(s) the relative strengths of all PASs decrease. However, any factor that would increase the window of opportunity for the CPA factors to recognize pPAS would lead to a shift toward the pPAS, such as RNAPII pausing, slowing RNAPII elongation, or increasing the distance between pPAS and dPAS. Even though there is still no evidence of this model in plants, it is an interesting putative scenario to see the light as a regulator of CPA factors abundance and PAS selection. Interestingly, we found that the distance between two APA sites is important for both types of APA isoform regulation, with 3’UTR lengthening and those with 3’UTR shortening (Suppl. Fig. 6).

As 3’UTRs are very rich in regulatory elements, the physiological consequences of the changes in APA decisions in response to light could be wide and affect many pathways. In particular, the gene with the event chosen as output, *HTA9*, encodes a histone H2A variant (H2A.Z). This variant has been associated with environmental responses to temperature and stress (45, 46). On the other hand, it was shown that H2A.Z-containing nucleosomes wrap DNA more tightly than canonical H2A nucleosomes, which may affect RNAPII elongation and, in turn, AS (7). In any case, further work will be necessary to better understand the biological roles of the novel link between plant APA and light reported here.

## Materials and methods

### Plant material, growth conditions and drug treatments

The *Arabidopsis* Columbia ecotype (Col 0) and Landsberg erecta (Ler) were used as wild type, according to the mutants assessed. Seeds were stratified for three days in the dark at 4°C and then germinated on Murashige and Skoog 0.5x (MS) medium containing 1% (w/v) agar. Standard treatment protocol: *A. thaliana* seedlings were grown in Petri dishes with MS medium at a constant temperature of 22°C under constant white light provided by fluorescent tubes with an intensity of irradiance between 70 and 100 μmol/m^2^sec (20 seeds per dish) for a period of 2 weeks and then transferred for 48hr to darkness. After this period, seedlings were kept in the dark or transferred to the light for 6 hours. At the end of this light/dark treatment, seedlings were harvested in liquid nitrogen. For all pharmacological treatments, drugs were added 1 hour prior to the light/dark treatment. For the DCMU [3-(3,4-dichlophenyl)-1,1-dimethylurea; Sigma] treatment seedlings were transferred to 6-wells plates and incubated in 20 µM DCMU. For the sucrose treatment, plants on agar plates were covered with 10 mL of 100 mM sucrose or sorbitol, used as osmotic control. Vacuum was applied for five minutes to facilitate drug uptake. For the DNP (2,4-dinitrophenol; Sigma) treatment, the drug was added up to 20 µM, using ethanol was as vehicle control.

### 3′READS and PAS identification

Total RNA extraction of seedlings was carried out by using the RNeasy Plant Mini Kit (Qiagen) following manufacturer’s instructions. The 3′ region extraction and deep sequencing (3′READS) method were described previously (11) Libraries were sequenced on an llumina HiSeq machine (2×150 bases) at Admera Health (New Jersey, USA). 3’READS data were analyzed as previously described (17, 47, 48). Briefly, after adapter sequence removal, reads were mapped to the *Arabidopsis thaliana* reference genome (TAIR10) by using the Bowtie2 program (49). Sequences with a mapping quality (MAPQ) score ≥ 10 and ≥ 2 nongenomic Ts at the 5′ end after alignment were considered as PAS-supporting (PASS) reads and were used for subsequence analysis. Identified cleavage sites within 24 nt from one another were clustered into PAS clusters (50). 3’READS data statistics are shown in Suppl. Table 1. Gene annotation was based on the TAIR10 and the Ensembl databases.

### Analysis of APA isoforms

Analysis of 3’UTR APA was based on the top two most expressed APA isoforms of a gene, which were named proximal and distal PAS isoforms. To eliminate spurious APA isoforms, we further required that the number of PASS reads for the minor APA isoform (the second most expressed) to be above 5% of all isoforms combined. For 3’UTR APA analysis, only the APA sites in the last exon were used. The relative expression (RE) of two PAS isoforms, e.g., pPAS and dPAS, was calculated by log2(RPM) of dPAS vs. pPAS, where RPM was reads per million PASS reads. Relative expression difference (RED) of two isoforms in two comparing samples was based on the difference in RE for the two isoforms between the two samples. DEXSeq was used to derive statistically significant APA events (FDR < 0.05) (51).

### Gene expression analysis

The DESeq method (R Bioconductor) (52)was used to analyze gene expression changes. Significantly regulated genes are those with adjusted *p* < 0.05 and fold change > 1.2. PAS reads of each gene were summed to represent gene expression.

### PAS motif analysis

The PROBE program was used to examine sequence motifs around the PASs (53). The genomic region surrounding each PAS was divided into four subregions: –60 to –31 nt, –30 to –1 nt, +1 to +30 nt, and +31 to +60 nt. The observed frequency of each k-mer in a subregion was enumerated and compared with the expected frequency based on randomized sequences of the region. Randomization was carried out by using the first-order Markov chain model (53). The enrichment score (*Z*-score) was calculated based on the difference between the observed and expected frequencies. The Fisher’s exact test was used to determine significance.

### Gene Ontology analysis

The GOstats hypergeometric test (R Bioconductor) was used to test for significant association of genes with gene ontology (GO) terms. GO annotation for *Arabidopsis thaliana* was obtained from org.At.tair.db (R Bioconductor). GO terms associated with more than 1,000 genes were considered too generic and were discard. To remove redundant GO terms, each reported GO term was required to have at least 25% of the genes that were not associated with another term with a more significant *p*-value.

### APA analysis by RT-qPCR

Seedlings were grown following specifications given in Plant material, growth conditions and drug treatments. Samples were harvested and total RNA was purified using TRIzol (Invitrogen). 500 ng of RNA were further used to synthesize cDNA with MMLV-RT enzyme (Invitrogen) and oligo-dT as primer following the manufacturer’s instructions. Synthesized cDNAs were amplified with 1.5 U of Taq DNA polymerase (Invitrogen) and SYBR Green (Roche) using the Eppendorf Mastercycler Realplex. Primer sequences for qRT-PCR are available in Suppl. Table 3. RT-qPCR experiments were quantified with n ≥ 3, where n = about 25-30 *Arabidopsis* seedlings growing in one Petri dish. Changes considered significant show differences with a p value < 0.05 (two-tailed Student’s t test).

## Acknowledgments

We thank E. Martín, V. Roselló and V. Buggiano for technical assistance. This work was supported by grants from the Agencia Nacional de Promoción Científica y Tecnológica of Argentina (PICT-2019-862), the Universidad de Buenos Aires (UBACYT 20020170100046BA) and the Lounsbery Foundation (USA) to ARK and NIH grants GM084089 and GM129069 to BT. M.A.G-H, E.P. and A.R.K. are career investigators and M.G.K. received a fellowship from the Consejo Nacional de Investigaciones Científicas y Técnicas of Argentina (CONICET).

## References

1. J. Yang, Y. Cao, L. Ma. Co-transcriptional RNA processing in plants: exploring from the perspective of polyadenylation. Int J. Mol. Sci. 22, 3300 (2021).

2. B. Tian, J. L. Manley. Alternative polyadenylation of mRNA precursors. Nat. Rev. Mol. Cell Biol. 18, 18–30 (2017).

3. M. Cyrek et al. Seed dormancy in Arabidopsis is controlled by alternative polyadenylation of DOG1. Plant Physiol. 170, 947–955 (2016).

4. F. Liu, S. Marquardt, C. Lister, S. Swiezewski, C. Dean. Targeted 3’ processing of antisense transcripts triggers Arabidopsis FLC chromatin silencing. Science 327, 94–97 (2010).

5. C. Xu, et al. R-loop resolution promotes co-transcriptional chromatin silencing. Nat. Commun. 12, 1790 (2021).

6. E. Petrillo et al. A chloroplast retrograde signal regulates nuclear alternative splicing. Science 344, 427–430 (2014).

7. M. A. Godoy Herz et al. Light regulates plant alternative splicing through the control of transcriptional elongation. Mol. Cell 73, 1066–1074 (2019).

8. A. R. Kornblihtt et al. Alternative splicing: a pivotal step between eukaryotic transcription and translation. Nat. Rev. Mol. Cell Biol. 14, 153–165 (2013).

9. T. Saldi, M. A. Cortazar, R. M. Sheridan, D. L. Bentley. Coupling of RNA polymerase II transcription elongation with pre-mRNA splicing. J. Mol. Biol. 428, 69–81 (2016).

10. S. Riegler et al. Light regulates alternative splicing outcomes via the TOR kinase pathway. Cell Rep. 36, 109676 (2021).

11. D. Zheng, X. Liu, B. Tian. 3′READS+, a sensitive and accurate method for 3′ end sequencing of polyadenylated RNA. RNA 22, 1631–1639 (2016).

12. A. Sherstnev et al. Direct sequencing of Arabidopsis thaliana RNA reveals patterns of cleavage and polyadenylation. Nat. Struct. Mol. Biol. 8, 845–852 (2012).

13. S. Zhu et al. PlantAPAdb: A Comprehensive Database for Alternative Polyadenylation Sites in Plants. Plant Physiol. 182, 228–242 (2020).

14. X. Liu et al. Comparative analysis of alternative polyadenylation in S. cerevisiae and S. pombe. Genome Res. 27, 1685–1695 (2017).

15. X. Liu et al. Transcription elongation rate has a tissue-specific impact on alternative cleavage and polyadenylation in Drosophila melanogaster. RNA 23, 1807–1816 (2017).

16. C. H. Jan, R. C. Friedman, J. G. Ruby, D. P. Bartel. Formation, regulation and evolution of Caenorhabditis elegans 3’UTRs. Nature 469, 97–101 (2011).

17. M. Hoque et al. Analysis of alternative cleavage and polyadenylation by 3’ region extraction and deep sequencing. Nat. Methods. 10, 133–139 (2013).

18. A. Nagatani, J.W. Reed, J. Chory. Isolation and initial characterization of Arabidopsis mutants that are deficient in phytochrome A. Plant Physiol. 102, 269–277 (1993).

19. M. Koornneef, E. Rolf, C.J.P. Spruit. Genetic control of light-inhibited hypocotyl elongation in Arabidopsis thaliana (L.) Heynh. Z. Pflanzenphysiol. 100, 147–160 (1980).

20. M. Koornneef, C.J. Hanhart, J.H. van der Veen. A genetic and physiological analysis of late flowering mutants in Arabidopsis thaliana. Mol. Genet. Genom. 229, 57–66 (1991).

21. M. Ruckle, L. Burgoon, L. Lawrence, C. Sinkler, R. Larkin. Plastids are major regulators of light signaling in Arabidopsis. Plant Physiol. 159, 366–390 (2012).

22. A. Khandelwal, T. Elvitigala, B. Ghosh, R. S. Quatrano. Arabidopsis transcriptome reveals control circuits regulating redox homeostasis and the role of an AP2 transcription factor. Plant Physiol. 148, 2050–2058 (2008).

23. K. Wippel, N. Sauer. Arabidopsis SUC1 loads the phloem in suc2 mutants when expressed from the SUC2 promoter. J. Exp. Bot. 63, 669–679 (2012).

24. J. Wind, S. Smeekens, J. Hanson. Sucrose: metabolite and signaling molecule. Phytochem. 71, 1610–1614 (2010).

25. Y. Xiong et al. Glucose-TOR signaling reprograms the transcriptome and activates meristems. Nature 496, 181–186 (2013).

26. D. Liu et al. The mitochondrial uncoupler DNP triggers brain cell mTOR signaling network reprogramming and CREB pathway up-regulation. J. Neurochem. 134, 677–692 (2015).

27. R. N. Fish, C. M. Kane. Promoting elongation with transcript cleavage stimulatory factors. Biochim. Biophys. Acta 1577, 287–307 (2002).

28. J. Dolata et al. NTR1 is required for transcription elongation checkpoints at alternative exons in Arabidopsis. EMBO J. 34, 544–58. (2015).

29. S. Sigurdsson, A. B. Dirac-Svejstrup, J. Q Svejstrup. Evidence that transcript cleavage is essential for RNA polymerase II transcription and cell viability. Mol. Cell 38, 202–210 (2010).

30. G. Dujardin et al. How slow RNA polymerase II elongation favors alternative exon skipping. Mol. Cell 54, 683–690 (2014).

31. L. E. Marasco et al. Counteracting chromatin effects of a splicing-correcting antisense oligonucleotide improves its therapeutic efficacy in spinal muscular atrophy. Cell 185, 2057–2070 (2022).

32. A. V. Klepikova, A. S. Kasianov, E. S Gerasimov, M. D. Logacheva, A. A. Penin. A high resolution map of the Arabidopsis thaliana developmental transcriptome based on RNA-seq profiling. Plant J. 88, 1058–1070 (2016).

33. Q. Zhang, B. Tian. The emerging theme of 3’UTR mRNA isoform regulation in reprogramming of cell metabolism. Biochem. Soc. Trans. 28, 1111–1119 (2023).

34. K. P. Forbes, B. Addepalli, A. G. Hunt. An Arabidopsis Fip1 homolog interacts with RNA and provides conceptual links with a number of other polyadenylation factor subunits. J. Biol. Chem. 281, 176–186 (2006).

35. R. Q. Xu et al. The 73 kDa subunit of the cleavage and polyadenylation specificity factor (CPSF) complex affects reproductive development in Arabidopsis. Plant Mol. Biol. 61, 799–815 (2006).

36. S. A. Bell, A. G. Hunt. The Arabidopsis ortholog of the 77 kDa subunit of the cleavage stimulatory factor (AtCSTF-77) involved in mRNA polyadenylation is an RNA-binding protein. FEBS Lett. 584, 1449–1454 (2010).

37. P. E. Thomas et al. Genome-wide control of polyadenylation site choice by CPSF30 in Arabidopsis. Plant Cell 24, 4376–4388 (2012).

38. G. Martin, A. R. Gruber, W. Keller, M. Zavolan. M. Genome-wide analysis of pre-mRNA 3’ end processing reveals a decisive role of human cleavage factor I in the regulation of 3’ UTR length. Cell Rep. 1, 753–763 (2012).

39. T. Kubo, T. Wada, Y. Yamaguchi, A. Shimizu, H. Handa. Knock-down of 25 kDa subunit of cleavage factor Im in hela cells alters alternative polyadenylation within 3’-UTRs. Nucleic Acids Res. 34, 6264–6271 (2006).

40. C. P. Masamha et al. CFIm25 links alternative polyadenylation to glioblastoma tumour suppression. Nature 510, 412–416 (2014).

41. M. Jenal et al. The poly(A)-binding protein nuclear 1 suppresses alternative cleavage and polyadenylation sites. Cell 149, 538–553 (2012).

42. C. Yao et al. Transcriptome-wide analyses of CSTF64-RNA interactions in global regulation of mRNA alternative polyadenylation. Proc. Natl. Acad. Sci. U.S.A. 109, 18773– 18778 (2012).

43. W. Li et al. Systematic profiling of poly(A)+ transcripts modulated by core 3’ end processing and splicing factors reveals regulatory rules of alternative cleavage and polyadenylation. PLoS Genet. 11: e1005166 (2015).

44. P. Tang, Y. Zhou. Alternative polyadenylation regulation: insights from sequential polyadenylation. Transcription 13, 89–95 (2022).

45. S. V. Kumar, P. A. Wigge. H2A.Z-containing nucleosomes mediate the thermosensory response in Arabidopsis. Cell 140, 136–47 (2010).

46. W. Sura et al. Dual Role of the Histone Variant H2A.Z in Transcriptional regulation of stress-response genes. Plant Cell. 29, 791–807 (2017).

47. D. Zheng, B. Tian. Polyadenylation site-based analysis of transcript expression by 3’READS. Methods Mol. Biol. 1648, 65–77 (2017).

48. D. Zheng et al. Cellular stress alters 3’UTR landscape through alternative polyadenylation and isoform-specific degradation. Nat. Commun. 9, article number 2268 (2018).

49. B. Langmead, S. L. Salzberg. Fast gapped-read alignment with Bowtie 2. Nat. Methods 9, 357–359 (2012).

50. B. Tian, J. Hu, H. Zhang, C. S. Lutz. A large-scale analysis of mRNA polyadenylation of human and mouse genes. Nucleic. Acids Res. 33 201–212 (2005).

51. S. Anders, A. Reyes, W. Huber. Detecting differential usage of exons from RNA-seq data. Genome Res. 22, 2008–2017 (2012).

52. S. Anders, W. Huber. Differential expression analysis for sequence count data. Genome Biol. 11, R106 (2010).

53. J. Hu, C. S. Lutz, J. Wilusz, B. Tian. Bioinformatic identification of candidate cis-regulatory elements involved in human mRNA polyadenylation. RNA 10, 1485–1493. (2005).

